# Microneedle delivery of stem cell derived retinal pigment epithelial cells

**DOI:** 10.1101/2025.04.29.651158

**Authors:** Jared Ching, Hinako Ichikawa, Kitahata Shohei, Liam Morrow, Kazuaki Kadonosono

## Abstract

Retinal pigment epithelium (RPE) transplantation surgery has shown potential benefits in animal models and early clinical trials. However, current implantation techniques are limited by the need for large retinotomies when cell bearing scaffolds are used or cellular reflux in the case of cell suspension transplantation with micro-cannulae. Here we demonstrate that it is feasible to pass primary and stem cell derived RPE cells through microneedles with a bore as small as 30μm. In vitro, there is no immediate loss of cell viability and this remains true throughout 14 days of live cell viability monitoring. Further, when the RPE cells are cultured over 14 days, they attach within 24 hours and become confluent, with normal morphological appearances. The formation of tight junctions is unimpeded with the typical polygonal pattern observed. The effects of shear stress due to passage of the cells through a small lumen were not evident in cultured cells labelled for the cytoskeletal protein F-actin. RPE cell VEGF secretion is unaffected by microneedle size in primary RPE and iPSC RPE cells. Ex vivo surgical implantation demonstrates that the technique is feasible, RPE cells engraft in the outer retina and we show that vitrectomy is not required to successfully transplant cells. Herein, we provide the first evidence of surgical microneedle cell delivery, which may have future applications more broadly within ophthalmic microsurgery and minimally invasive delivery strategies.

**Graphical summary:** 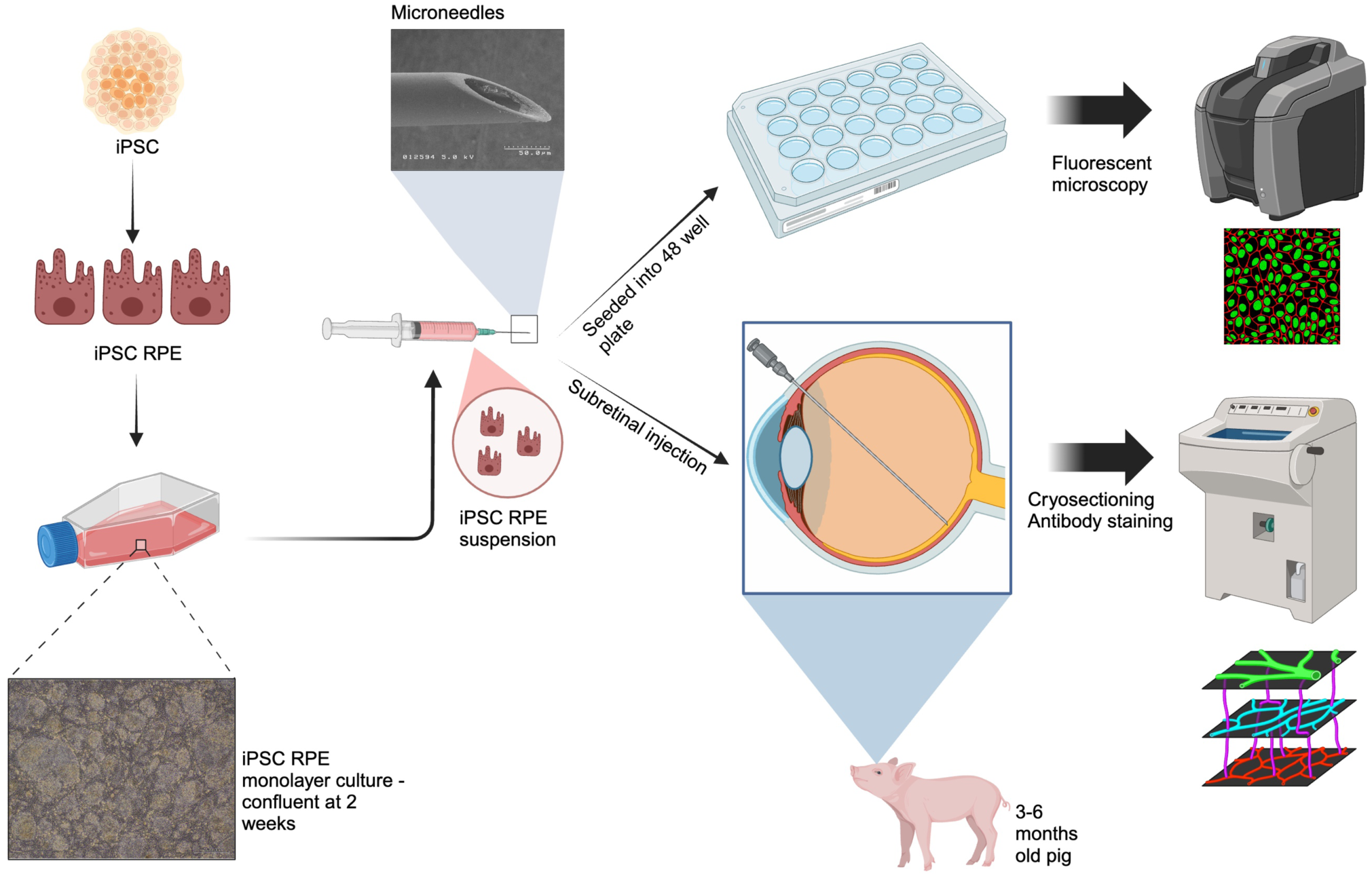

Induced pluripotent stem cells (iPSC) are differentiated into retinal pigment epithelial cells (iPSC RPE) and cultured for 14 days until confluent. A single cell suspension of iPSC RPE is then injected using an Alcon Constellation Vision System machine through different microneedles into a 48 cell culture plate or in the subretinal space of an ex vivo pig eye 3-6 months in age. Cells and tissues were processed and stained by using various antibodies to assess viability, morphology, function and anatomical localization in the subretinal space to confirm the feasibility of microneedle cell delivery.

## Introduction

Degenerative retinal diseases such as age-related macular degeneration and retinitis pigmentosa, collectively represent significant causes of irreversible sight loss globally.^1^ Currently, there is a paucity of treatment options where cell therapy with stem cell derived retinal pigment epithelium cells (RPE) has emerged with promising pre-clinical data. ^2,3^ The two categories of RPE transplantation techniques are scaffold-based and scaffold-free.^4^ Both techniques have their advantages and limitations. Scaffold-based methods tend to require sizable retinotomies to access the subretinal space, whereas scaffold-free methods are less reliant this, particularly when considering cell suspension subretinal delivery using a small gauge cannula.^5^ However, despite the small retinotomy created by a 38 gauge subretinal cannula when delivering RPE suspensions, transplanted RPE cells are thought to reflux through this wound into the vitreous cavity and come into contact with the inner retina, leading to epiretinal membrane formation.^6^ Efforts to reduce this phenomenon have included the development of a gelatin hydrolysate cell suspension vehicle that has been shown to reduce RPE cell reflux in an in vitro retinal bleb model.^7^ Further, RPE strips have been developed as a compromise between a RPE scaffold-free sheet and cell suspension.^8,9^

Retinal microneedles, defined as having at least one dimension between 1 to 999µm, have been developed in recent times to facilitate reliable, safe and effective means to cannulate retinal vasculature with the goal of treating vaso-occlusive diseases.^10^ Patients treated with vitrectomy surgery and microneedle endovascular cannulation, which can be augmented with recombinant tissue plasminogen activator, have been shown to potentially benefit from anatomical, angiographic and functional improvements.^11,12^ Such microneedles are available commercially and can be used seamlessly in conjunction with vitrectomy systems such as the Alcon Constellation Vision System and DORC EVA NEXUS. The microneedle developed by Tochigi Seiko, Tochigi, Japan, has an outer diameter of 50μm and estimated inner lumen diameter of 35μm.^10^ In comparison, the most commonly used and least Invasive subretinal cannula (Polytip 25/38G, MedOne) used in RPE cell suspensions delivery has an outer diameter of 38G (ca. 111μm) and inner lumen of 41G (78μm).^6,13,14^ The microneedle therefore has a 2.22 times smaller inner and outer diameter that may result in smaller retinotomy.

To our knowledge, the extent of minimally invasive surgical delivery of cell therapy has yet to be fully explored. We sought to investigate whether microneedle cell delivery is feasible using in vitro and ex vivo methods.

## Methods

### Microscopy of microneedles

A Leica S9D stereomicroscope (Leica Microsystems GmbH, Wetzler, Germany) used to measure the needle tip were undertaken following calibration of the scale at a fixed zoom of 10X. All images were undertaken using a Flexacam C3 microscope camera (Leica Microsystems GmbH, Wetzler, Germany). All images were exported into ImageJ and analysed by zooming maximally to take measurements of the outer diameter, wall thickness and inner diameter.

### Electron microscopy of microneedles

All microneedles were stabilised on a stage using carbon tape and scanning electron microscopy was undertaken using a Keyence VE-8800. All imagees were undertaken using accelerating voltages at 0.8kV or 2.0kV.

### Shear stress calculation

As the cells pass through the needle, they experience shear stress imposed by the fluid. To estimate the flow conditions, we consider Reynold’s number

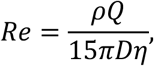

where 𝜌, Q, D, and 𝜂 are the density, flow rate, needle diameter, and dynamic viscosity of the fluid. Taking 𝜂 = 0.001 𝑃𝑎 𝑠 and 𝜌 = 997𝑘𝑔/𝑚^!^, we find that the Reynolds number is sufficiently small over the values of D and Q considered in this work such that we assume that the behavior of the fluid is described by Poiseuille flow. As such, the velocity profile and shear stress of the fluid are parabolic and linear, respectively, as illustrated in Fig. 1. This figure shows that the maximum shear stress will be on the walls of the needle, which also corresponds to where the fluid is stationary.

**Figure 1:**
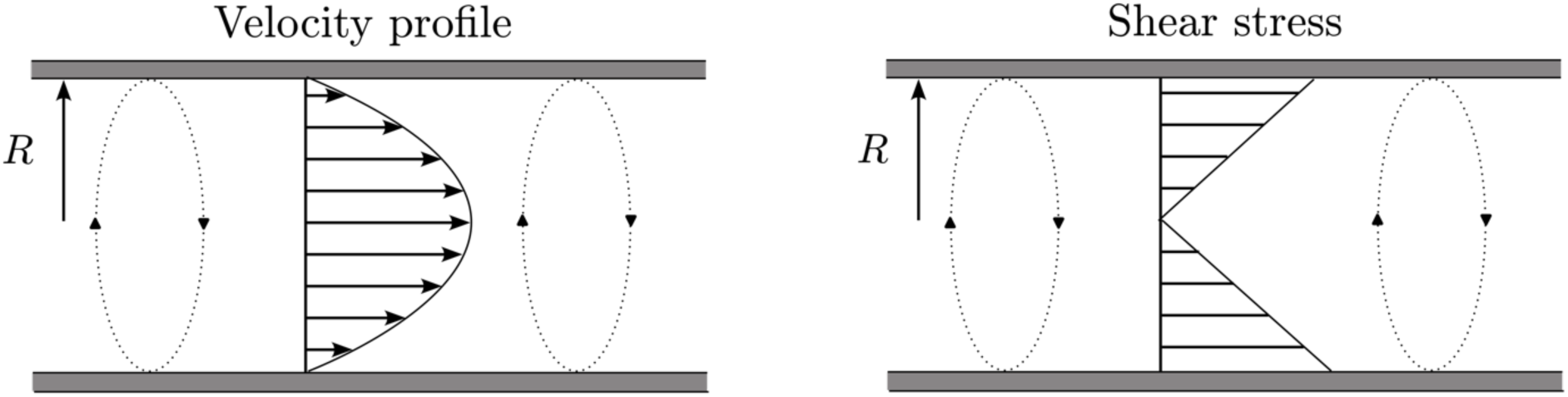
Velocity profile and shear stress of fluid are parabolic and linear during passage through the bore of a microneedle.

**Figure 1:**
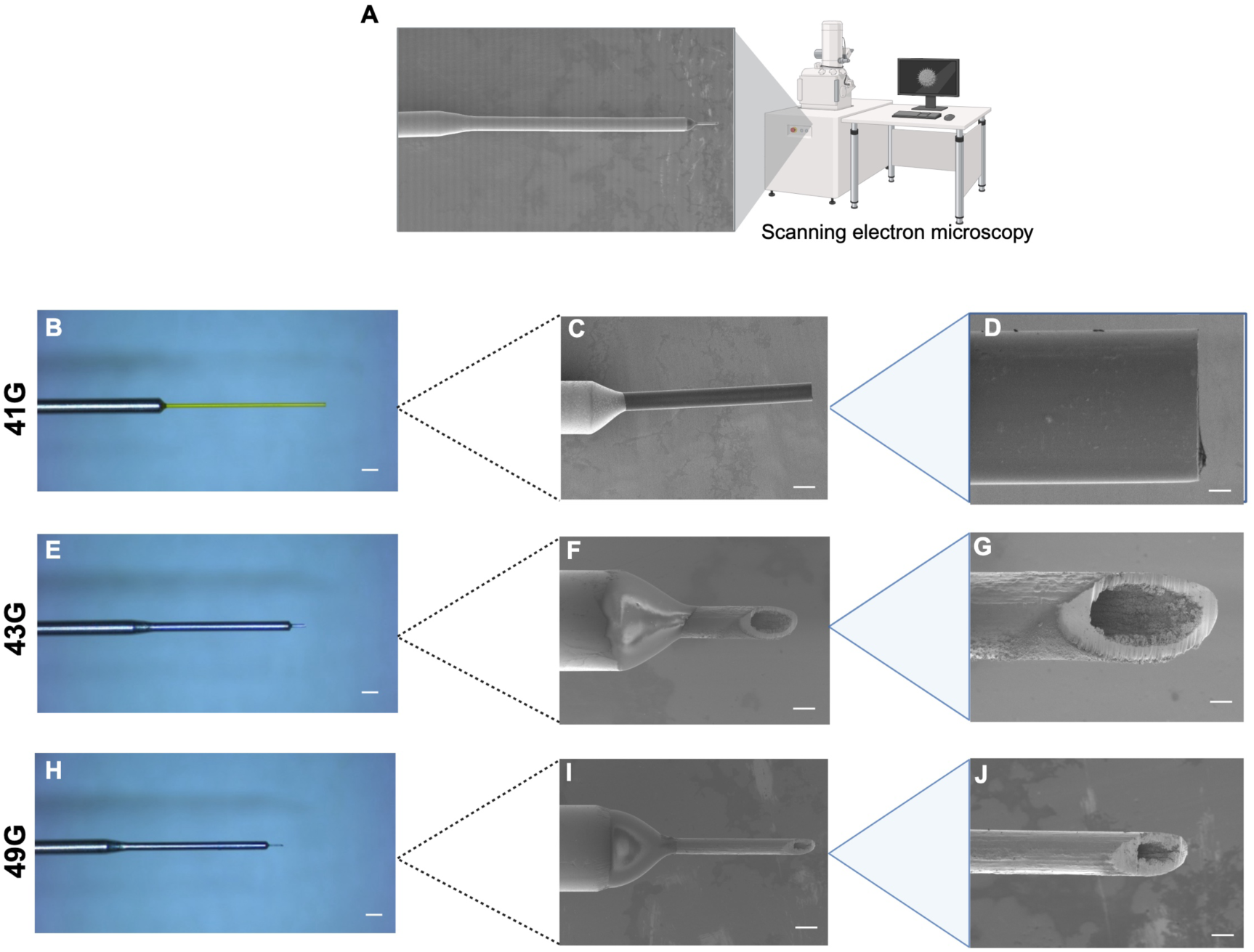
A) Schematic of scanning electron microscopy (SEM) images of microneedles. Colour photomicrographs of the microneedles used in the present study, including the 25G/38G Polytip with an internal diameter of 41G and 5mm length(B), microneedle with an internal diameter of 43G (E), and microneedle with an internal diameter of 49G (H), where scale bars indicate 500µm. SEM at 50x magnification of a the 41G Polytip with a 3mm tip (used instead of the 5mm for imaging purposes only) where the scale bar represents 200µm (C). SEM of 43G and 49G microneedles at 150x magnification (F & I), where the scale bar represents 6.66µm. The microneedles examined at 400x magnification under SEM (D, G, J), where the scale bar represents 25µm.

Given the fluid velocity is fastest along the centreline of the tube, it is expected that the cells tend to will migrate towards centre of the needle where the shear stress is at its smallest. As such, we approximate the shear stress imposed by the fluid on the cells by taking a cross-sectional average to give

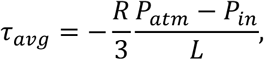

where 𝑃*_atm_*, 𝑃*_in_*, and 𝐿 are atmospheric pressure, the input pressure and the length of the needle. We note that this value of average shear stress corresponds to 2/3 the value of the maximum shear stress on the walls of the needle.

### Primary human RPE cell culture and Human iPSC-derived RPE cell culture and suspension preparation

The primary human RPE cells (#00194987) were purchased from Lonza. RPE cells were differentiated from the established human iPSC line 253G1^15^ (RIKEN Bioresource Center, Tokyo, Japan) using a previously described protocol.^16^ RPE cells were differentiated from the established human iPSC line 253G1^15^ (RIKEN Bioresource Center, Tokyo, Japan) using a previously described protocol.^16^ Briefly, iPSC. Line 253G1 was cultured StemFit AK03 (Ajinomoto Co., Tokyo, Japan) on laminin-511 E8 (iMatrix-511, Nippi Inc., Tokyo, Japan)-coated plates. Differentiation into RPE cells was undertaken as described previously^16^ by using a polymerized mixture of 7:2:1 ratio of acid solubilized porcine tendon collagen type I-A (3 mg/ml, Cellmatrix; Nitta Gelatin), 5X concentrated Dulbecco’s modified Eagle’s medium (DMEM; Nitta Gelatin), and reconstitution buffer (NaOH 50 mM, NaHCO_3_ 260 mM, HEPES 200 mM; Nitta Gelatin). The iPSC-RPE were cultured in serum-free retinal medium supplemented with 10 ng/ml basic fibroblast growth factor and SB431542 (0.5 μM) after reaching confluence. Media was changed every 2-3 days and after 4 weeks when typical RPE cobblestone morphology was evident, the iPSC-RPE cells were dissociated with trypsin and seeded into Nunc low cell surface binding flasks (Life Technologies, Carlsbad, CA, USA) with maintenance medium consisting of DMEM-Ham’s F12 basal medium (Sigma-Aldrich, St Louis, MO). The confluent colonies were then transferred to 12-well plates coated with CELLstart (Lifee Technologies, Carlsbad, CA) for 3 weeks before subculture and stored as passage 2 (P2) using CELLBANKER 1 plus (Nippon Zenyaku Kogyo Co., Fukushima, Japan). Cells were stored at 150 °C and thawed on CELLstart-coated plates and expanded as needed.

The primary human RPE and iPSC-RPE were cultured in serum-free retinal medium supplemented with 10 ng/ml basic fibroblast growth factor and SB431542 (0.5 μM), which were replaced every 2-3 days until the cells became confluent. Both RPE cells were dissociated with trypsin, suspended in culture medium with 5 μM Y27632, and used for the experiments.

### Surgical instrumentation

For all experiments, the Constellation Vision System (Alcon, Fort Worth, TX, USA) was used. For all experiments, 25 Gauge Vitrectomy packs were used with the Viscous Fluid Control Pack (BL7600, Alcon, Fort Worth, TX) that included a 10mL syringe, syringe plunger, syringe cap, tubing set with syringe coupler, 20G, 23G and 25G cannula. For all experiments a MicroDose™ Injection Kit (3275, MedOne, Saratosa, FL) was used in place of the 10mL syringe included in the Viscous Fluid Control Pack for more accurate fluid manipulation. The microneedles used included the 25G/38G Polytip® (MedOne, Saratosa, FL), Tochigi Seiko 0.11mm prototype microneedle (Tochigi Seiko Co. Ltd., Tochigi, Japan), and the Tochigi Seiko 0.05mm microneedle (Tochigi Seiko Co. Ltd., Tochigi, Japan). All experiments carried out in vitro utilized a HEPA filtered sterile hood alongside sterile surgical instruments.

In addition to the aforementioned equipment, further surgical instruments for ex vivo surgery included balanced salt solution, ophthalmic viscoelastic Opegan 0.6 (Santen Pharmaceutical Ltd., Osaka, Japan), posterior vitrectomy contact lens (Hoya Corporation, Shinjuku), 22.5 degree blades (MANI Inc., Tochigi, Japan), toothed forceps (Duckworth and Kent, Baldrock, UK), and curved scissors, spring scissors.

### Cell viability assessments

The standard trypan blue exclusion assay was performed using the automatic counting function of TC20™ Cell Counter (Biorad, Hercules, CA) and manual counting using disposable haemocytometers (Watson Biolab, San Diegao, CA) under bright field microscopy. The average of the two methods were used or where the automated counting failed, only the manual counts were included. SYTOX™ Green nucleic acid stain 5mM solution was used at a 1:1000 concentration to monitor cell death during cell culture on the same day as seeding, and after 1, 3, 7, and 14 days of culture. Images were transferred to ImageJ (National Health Institutes, Bathesda, USA), processed in black and white and the same threshold applied to all images before quantification of fluorescent dots.

### Immunocytochemistry

Cells were washed in PBS before fixing in 4% paraformaldehyde (Polysciences, Pennsylvania, USA) for 15 minutes at room temperature and pressure. A further 3 x 5 minute washes with PBS was performed before blocking (Blocking One, Nalacai Tesque, Kyoto, Japan) for 1 hour at room temperature and pressure. The primary antibody ZO-1 1:1000 (Thermo Fisher Scientific, Massachusetts, USA) was added and incubated at 4°C overnight before washing with PBS 3 x 5 minutes. The secondary antibody Alexafluor 488 (Thermo Fisher Scientific, Massachusetts, USA) was added at 1:1000 or phalloidin 594 (Thermo Fisher Scientific, Massachusetts, USA) with DAPI 1:100. The cells were washed with PBS 3 x 5 minutes once more before imaging.

### Microscopy

Fluorescent microscopy was carried out with a Kayence BZ-X710 Fluorescent Microscope (Kayence Tokyo Laboratory, Tokyo, Japan). Briefly, four channels including bright field, DAPI, GFP and TRITC were used to image all cell culture, retinal wholemount samples and retinal sections at magnifications of 4X, 10X and 20X. All image processing was undertaken using the Kayence Analysis software package and ImageJ.

### Ex vivo pig eye model

In order to confirm the feasibility of microneedle delivery of RPE cells to the subretinal space, we utilized an ex vivo pig eye model akin to Gatto et al.^17,18^ Pig eyes were harvested from freshly terminated animals 3-6 months old from abattoirs within. The vicinity. The eyes were delivered on ice with the extraocular muscles, conjunctiva, eyelids and tissues intact and arrived at the laboratory within 2-4 hours of termination. The pigs eyes were mounted onto a polystyrene phantom head and washed with double distilled water (ddH_2_O). Using the Constellation Vision System (Alcon, Texas, USA), a 25 gauge three port vitrectomy was performed on each pig eye. A vitrectomy contact lens (Hoya, Tokyo, Japan) was used as the posterior viewing system. A posterior vitreous detachment was performed and the posterior hyaloid membrane was peeled with end gripping forceps where necessary to clear posterior vitreous where necessary. The viscous fluid control (Alcon, Texas, USA) set was used with the included 20 gauge cannula to extract RPE cell suspensions into a 1mL MicroDose syringe (MedOne with or without fluorescent microbeads). The cannula was then exchanged for the needle of interest Polytip 25/38G (MedOne), Tochigi Seiko 0.11mm or Tochigo Seiko 0.05mm needles before the RPE suspension was injected. The extraction settings were set to a maximum of 650mmHg and the inject settings were set to a maximum of 4PSI for the Polyip 25/38G and Tochigo Seiko 0.11mm. Due to the slower flow rate found with the Tochigo Seiko 0.05mm microneedle, the injection pressures ranged between 4 to 12 PSI using the linear foot pedal control so that suspension media was seen gradually forming droplets at needle tip.

### Ex vivo imaging

Imaging was carried out using a Cirrus 5000 (OCT) System (Zeiss, Oberkochen, Germany). The pigs eyes, while mounted in the phantom head, were imaged. The cornea was hydrated with balanced salt solution and viscoelastic. The corneal epithelium was removed if causing impairment of visualizing the retinal structures. The scanning sequences used included the macula cube and 5 raster scans. The globes were rotated with in the simulated orbit as necessary to image the region of interest and the phantom head was manipulated and rotated as necessary to optimize imaging as needed.

### Histology

Following surgery, the enucleated pigs eyes were dismounted from the phantom head and an anterior chamber paracentesis was made at the limbus with a 22.5° angled blade (Mani Inc., Tochigi, Japan) followed by two further stab incisions at the pars plana to facilitate ingress of fixative. The globes were then placed in 4% paraformaldehyde phosphate buffer solution (Nacalai Tesque, Kyoto City, Japan) at 4°C overnight before immersion in 10%, 15% and 20% sucrose in PBS solution for 12 hours each at 4°C. Following this, the eyes were coated and immersed in OCT medium and frozen using a metal plate over dry ice. The globes were then placed within an embedding cup before cryosectioning at 10µm thickness using a Leica CM1950 Cryostat (Leica, Wetzler, Germany).

Sections were placed on a Super Frost coated glass slide (Matsunami Glass Ind. Ltd., Osaka, Japan), dried and stored at −20°C. Sections were washed with PBS permeabilized with Triton-X 0.1% for 1 hour at room temperature and pressure (RTP), blocked for 1 hour RTP and the primary antibodies added for 1 hour RTP. Slides were washed 3 times with PBS before the secondary antibodies and DAPI were added for 1 hour RTP before washing 3 times and mounting for imaging.

Haemotaxylin and eosin staining was performed using a standard protocol, briefly, sectioned tissues were immersed in double distilled water (ddH_2_0) before applying haemotoxylin to cover the tissues for 3 minutes. The slides were rinsed twice with ddH_2_0 before applying bluing reagent for 15 seconds. After rinsing with ddH_2_0, the slides were immersed in 100% ethanol and eosin was then applied. The slide was rinsed twice with 100% ethanal and then immersed in 100% ethanol before mounted with a cover slip.

### Statistical analysis

All statistical analysis was undertaken using GraphPad Prism 8.4.0 (455) for MacOS. Comparisons were analyzed using 1 way ANOVA with Tukey’s. post-hoc pairwise comparisons, where statistical significance was defined as p < 0.05.

## Results

### Microneedle internal diameter estimation

The majority of hypodermic needle sizes are most widely defined by gauge according the US Birmingham Wire Gauge or British Standard Wire Gauge along with the International Organization for Standardization (ISO) Color Coding.^19^ These systems describe gauge in terms of the outer diameter of the needle and are only defined in their metric measurements in millimeters up to 34 gauge. Therefore the inner diameter of needles smaller than this are generally measured or estimated by individual manufacturers, without any universal standards. Therefore, we undertook microscopy to estimate the inner diameter in relation to the outer diameter and needle wall thickness. The results of these measurements are summarized in Table 1.

**Table 1:** Photomicroscopy measurements of each needle and control with estimated internal gauge based on the ISO and British Standard Wire Gauge.

| Needle/Control | Internal diameter | Outer diameter | Wall thickness | Estimated internal gauge |
| --- | --- | --- | --- | --- |
| 25G/38G Polytip® | 0.08 | 0.11 | 0.01 | 41G |
| Tochigi Seiko 0.11mm prototype microneedle | 0.06 | 0.11 | 0.04 | 43G |
| Tochigi Seiko 0.05mm microneedle | 0.03 | 0.05 | 0.01 | 49G |
| Standard 200µL pipette tip (Control) | 0.50 | 0.95 | 0.19 | 25G |

### Shear Stress calculations

In order to understand the degree of shear stress cells being delivered as a cell suspension through microneedles, we use the equation described above. The values of the estimated average shear stress for each needle are given in Table 2. These results indicate that the despite having the smallest internal radius, the short 49G needles have lower average shear stress compared to the 43G needle due to their shorter length resulting in a smaller pressure drop. As expected the 34G and 30G needles have a lower average shear due to their larger internal radius.

**Table 2:** Table of average shear values calculated from the parameters of each needle.

| Needle type | Internal Radius ($\mu\text{m}$) | Length (cm) | Injection pressure (PSI) | Average shear ($\text{dyn}/\text{cm}^2$) |
| --- | --- | --- | --- | --- |
| 30G | 60 | 1.3 | 4 | 113 |
| 34G | 50 | 0.8 | 4 | 154 |
| 41G | 40 | 0.3 | 4 | 328 |
| 43G | 25 | 0.026 | 4 | 2360 |
| 49G | 16.25 | 0.004 | 4-12 | 999 |

### RPE cell viability is not significantly affected by microneedle gauge or cell density

In order to assess immediate cell viability following microneedle injection of cell suspensions of primary and iPSC derived RPE cells, the trypan blue exclusion assay was utilized with a full vitrectomy surgical set up adjacent to a laminar flow hood for sterile cell culture (Fig. 2A and 2B). Two different cell densities were used to assess whether this affected cell viability, including 6.0 x 10^4^ (Fig. 2C) and 2.5 x 10^5^ (Fig. 2D) based on previously reported cell suspension transplantation experiments and trials.^6–8^ Automated and manual counts were taken to determine the total and viable cells present in each of the conditions. We found that when injecting 6.0 x 10^4^ iPSC RPE cells a total of 1.42 x 10^5^ (1.61 x 10^5^), 1.03 x 10^5^ (standard deviation 5.51 x 10^4)^, 2.37 x 105 (2.20 x 10^5^), and 9.91 x 10^4^ (7.96 x 10^4^) for the control, 41G, 43G and 49G microneedles, respectively. Of these cells, 1.24 x 10^5^ (1.29 x 10^5^), 8.33 x 10^4^ (3.82 x 10^4^), 1.92 x 10^5^ (1.83 x 10^5^), 9.19 x 10^4^ (7.53 x 10^4^), were found to be viable, i.e. cells with no uptake of trypan blue, representing 87%, 81%, 81% and 83% cell viability, respectively. In comparison, when injecting a higher density used in both preclinical and clinical trials of 2.50 x 10^5^ iPSC RPE cells, a total of 6.73 x 10^5^ (1.29 x 10^5^), 4.72 x 10^5^ (3.82 x 10^4^), 2.88 x 10^5^ (1.83 x 10^5^), 2.59 x 10^5^ (7.53 x 10^4^). Of these cells, 6.22 x 10^5^ (1.29 x 10^5^), 4.54 x 10^5^ (3.82 x 10^4^), 2.88 x 10^5^ (1.83 x 10^5^), 2.59 x 10^5^ (7.53 x 10^4^), were found to be viable, representing 92%, 96%, 79% and 83% cell viability, respectively. No statistically significant difference between groups were found. This experiment was undertaken following validation that primary RPE cells can be injected with microneedles and demonstrate high levels of cell viability in the same conditions presented here (Supplementary Figure 1) .

**Figure 2:**
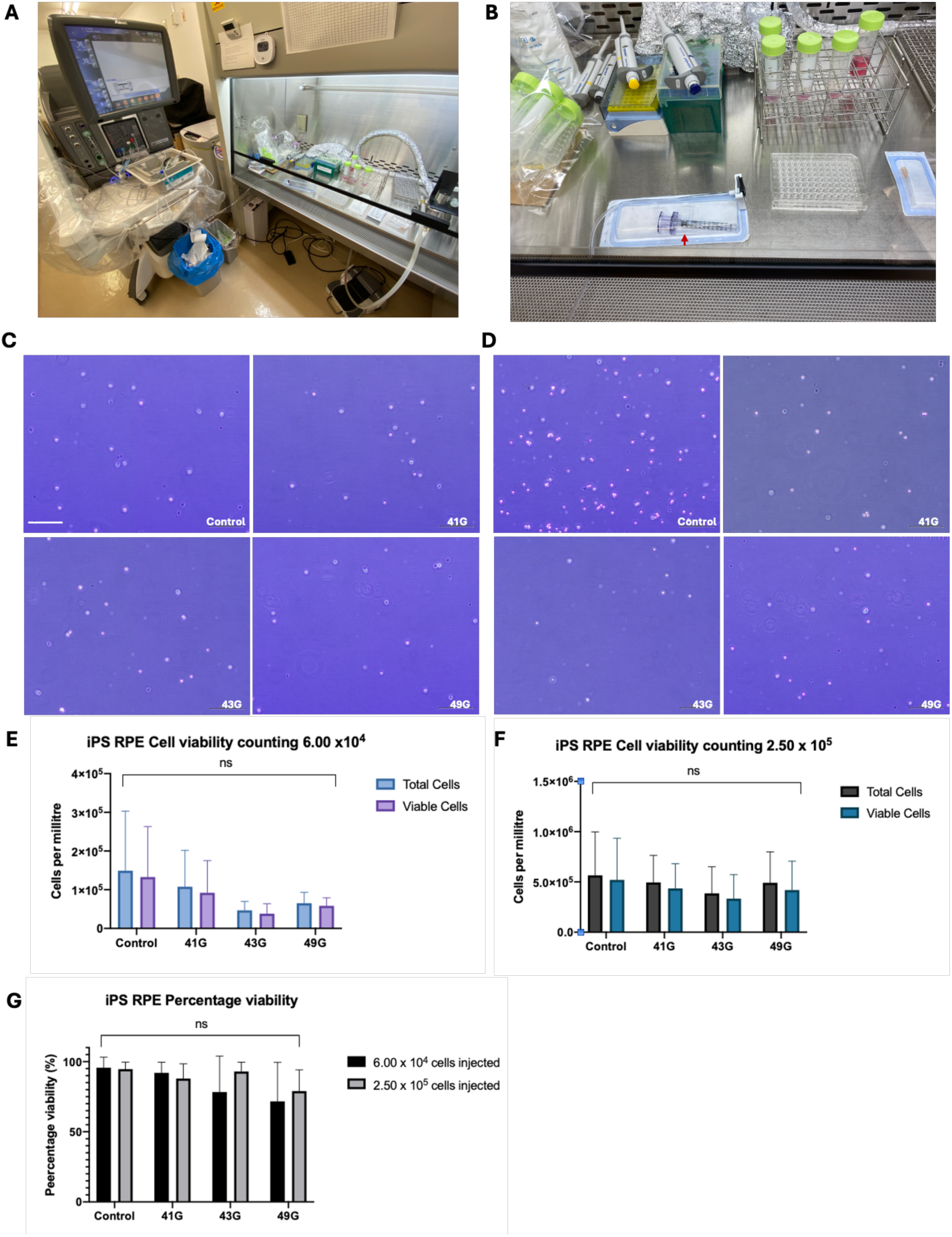
Cellularity and viability of iPSC RPE cells injected from microneedles. (A) Alcon Constellation Vision System set up adjacent to a laminar flow hood, used to inject cells through microneedles to investigate viability, morphology and function. (B) Sterile instruments used to culture RPE cells, including the VFC connector and MedOne Microdose 1mL syringe (red arrow). (C) Trypan blue exclusion assay imaged under bright field microscopy for iPSC RPE cells injected at a density of 6 x 10^4^ in 150µL, scale bar represents 500µm which all applies to all images in (C) and (D). (D) Trypan blue exclusion assay image under bright field microscopy for iPSC RPE cells injected at a density of 2.5 x 10^5^ in 150µL. (E) Total and viable cells injected from control (P200) and different microneedles as calculated from trypan blue exclusion assays using cell densities of 6 x 10^4^ in 150µL and (F) 2.5 x 10^5^ in 150µL. (G) Percentage cell viability for control and microneedles with different cell concentrations. abbreviations: ns = non-significant, G = gauge (lumen diameter).

In order to monitor cell death in vitro over time, we concurrently used SYTOX™ green nucleic acid stain immediately after injecting the cells into a 48 well cell culture plate and on days 1, 3, 7, and 14 quantified the number of dead cells. We found that there was no significant difference in the number of dead cells between cells injected with each of the microneedles when compared to the control (Fig. 3A). There was also no statistically significant change in the number of fluorescent dots between days 1, 3, and 7 in each condition. However, a significant dropout in signal was noted at day 14 in all conditions, with no evident increase in loss of cell viability.

**Figure 3:**
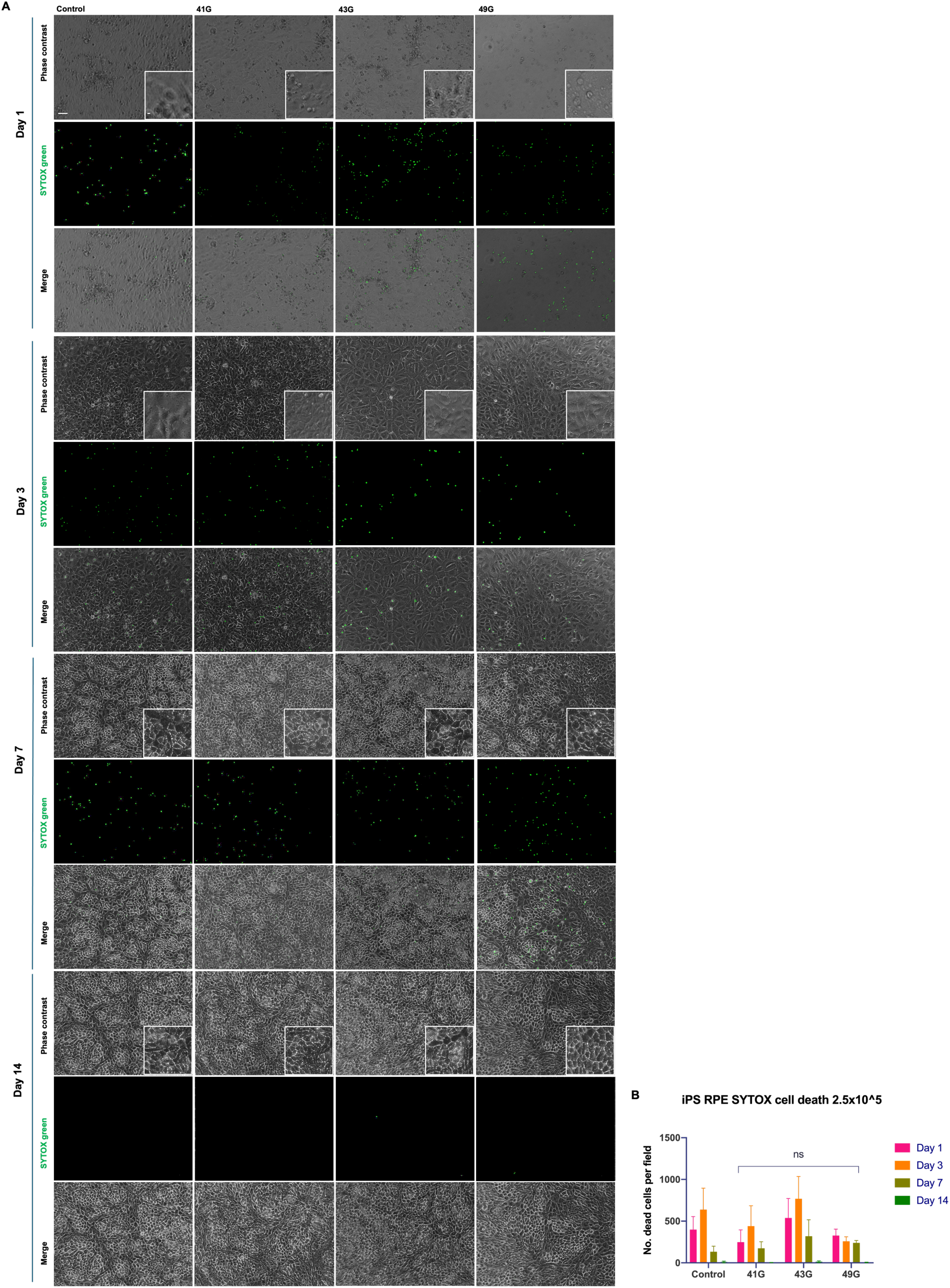
(A) Time course of primary RPE cells cultured after injection with control and microneedles. Brightfield microscopy at 10X and 20X zoom in the centre of each 48 well plate. (B) Graph shows corresponding cell death using SYTOX Green nuclear marker to quantify the number of dead cells per field. No statistical difference was found in SYTOX green staining when comparing to control at each time point. Scale bars in top left image represent 50µm and within the inset 10µm, which apply to all respective images in the panel.

### Cell morphology and cyotoskeleton is not adversely affected by microneedle gauge

The gross cell morphology of the iPSC RPE cells was assessed following microneedle injection and seeding of the cells into 48-well plates coated with laminin-511 E8 (iMatrix-511, Nippi Inc., Tokyo, Japan). Cells were cultured for a total of 14 days and imaged at day 0, 1, 3, 7 and 14 at 37°C, 5% CO_2_ and maintenance media changed every 2-3 days. High magnification phase contrast images of the cells at different time points appeared to show no difference with the controls, indicating that smaller microneedle gauges do not affect the ability of transplanted cells to proliferate (Fig. 3). Tight junction formation was assessed with zona occludens (ZO-1), along with DAPI. No difference was found in comparison to controls and the hexagonal typical conformation was present in all samples, indicating that microneedle gauge does not affect the ability of transplanted cells to form tight junctions (Fig. 4). Finally, it is known that shear stress can cause significant cytoskeletal changes to cells. F-actin was used to assess any cytoskeletal damage, however, no deviation from controls was seen or loss of lateral circumferential bundles (Fig. 5A). Further, we confirmed that cell viability was not adversely affected by needle gauge by comparing the total number of cells using DAPI staining, and dead cells, using SYTOX green. We found that percentage cell viability at day 14 was 98.37% (Standard deviation: ± 1.34%), 99.54% (± 0.39), 98.74% (± 1.14), 99.35% (± 0.45) (Fig. 5B & 5C), for the control, 41G, 43G, and 49G needles, respectively.

**Figure 4:**
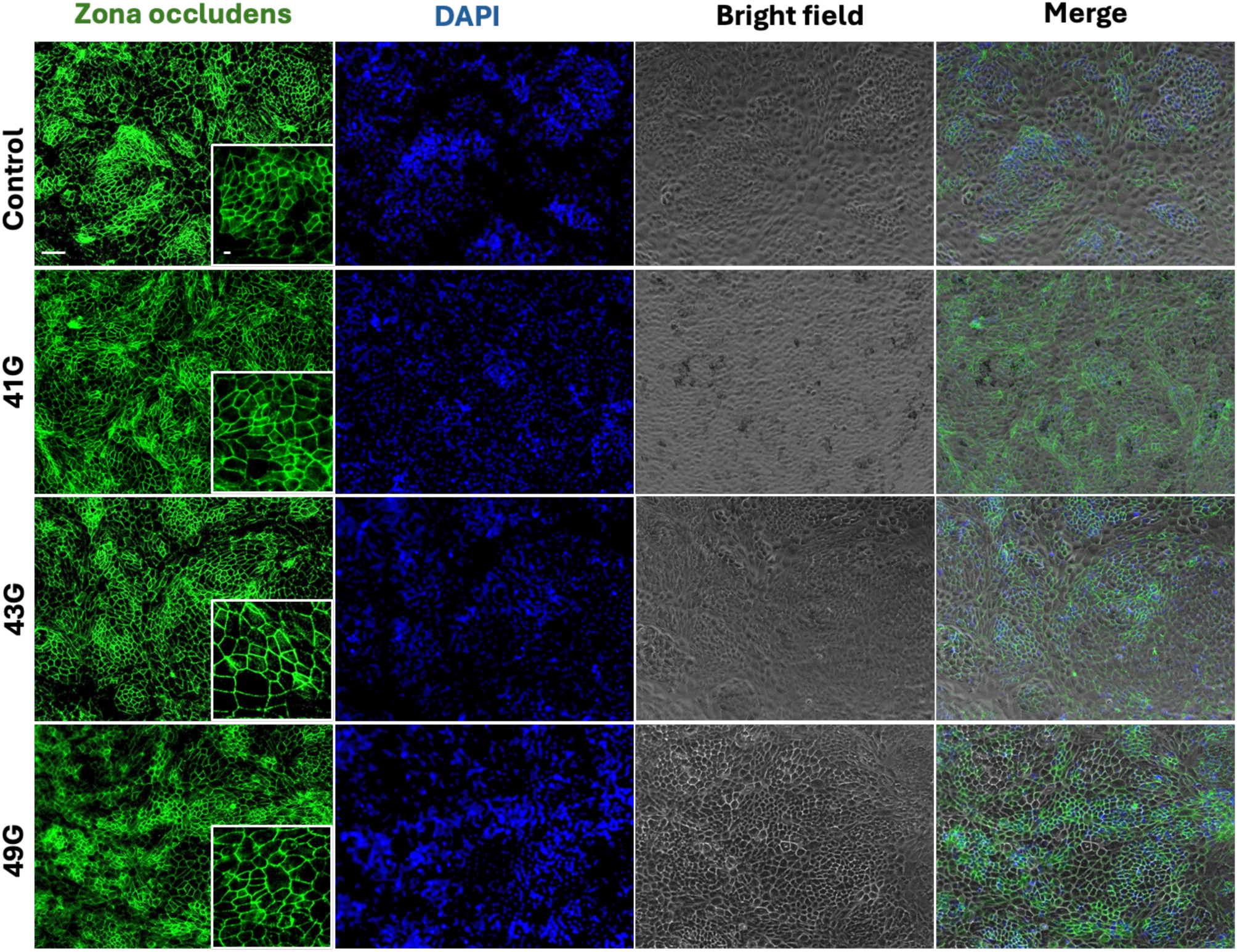
iPSC RPE cell morphology. Scale bars in top left image represent 50µm and within the inset 10µm, which apply to all respective images in the panel.

**Figure 5:**
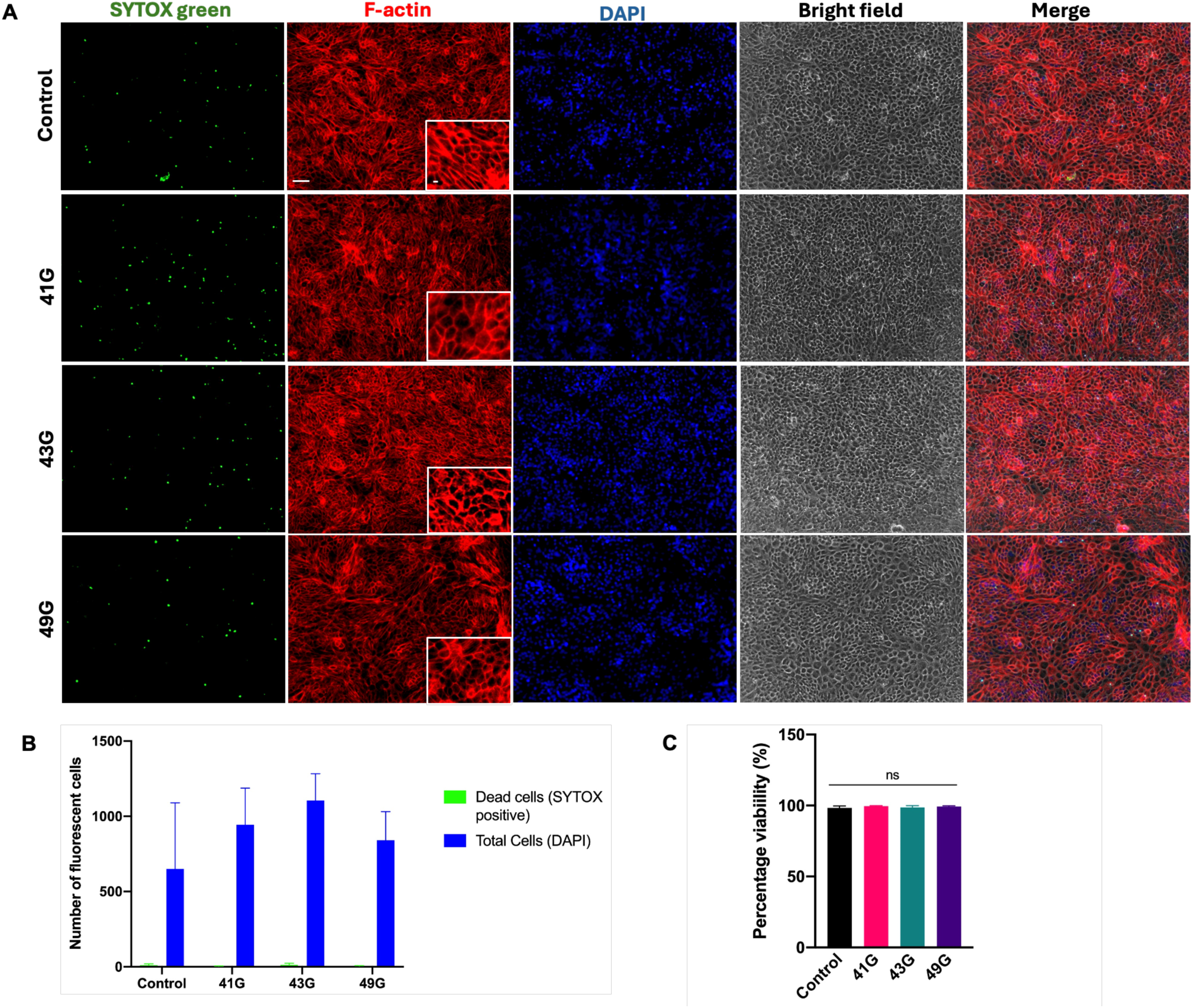
iPSC RPE viability and cytoskeletal status following in vitro seeding using microneedles attached to a vitrectomy machine. (A) F-actin distribution appears unaffected by needle gauge when compared to controls. Scale bars in the Control F-actin image represent 50µm and within the inset 10µm, which apply to all respective images in the panel. (B) Total cells indicated by counting the total number of DAPI positive cells and dead SYTOX green positive cells, with the percentage viability calculated accordingly (C).

### VEGF production is unaffected by smaller microneedle gauges

Function of the iPSC RPE cells was assessed by examining the trophic factor vascular endothelial growth factor (VEGF) by ELISA. We found that the smaller gauges of 43G and 49G was associated with less VEGF being detected on average, with the media of cells delivered by the 49G needle demonstrating a statistically significant lower concentration of VEGF (p = 0.04), Fig. 6A. However, when considering the viable cell count obtained from each needle (Fig. 6B), and then the amount of VEGF secreted per cell, the VEGF concentration appeared to be comparable within 1.0 x 10^-3^pg/mL between all conditions (Fig. 6C). Such variability was also seen with primary RPE cells, which demonstrated an opposing pattern of significantly higher VEGF concentrations being found for 41G and 43G microneedles compared to control (Supplementary Fig. S2A). However, when controlling for cell number (Fig. S2B), the VEGF concentration was also within 1.0 x 10^-3^pg/mL (Fig. S2C).

**Figure 6:**
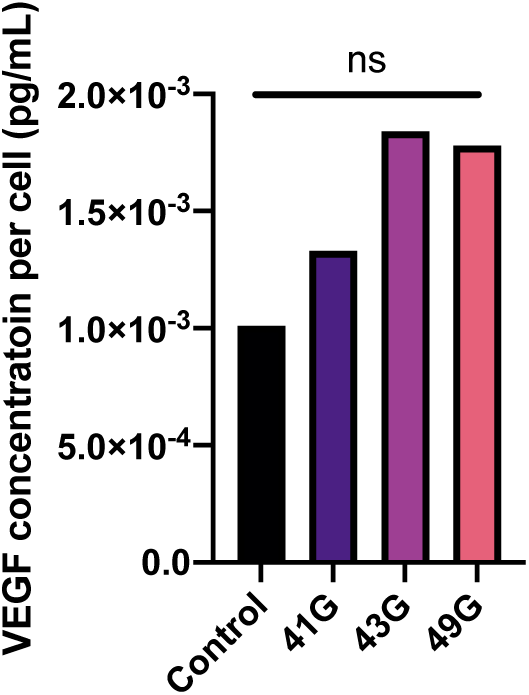
Normalised iPSC RPE VEGF secretion measured by ELISA. Controlling for cell number, the normalised amount of VEGF secreted per cell was within a range of 1.0 x 10^-3^pg/mL between all conditions with no significant difference between conditions.

### Stem cell derived RPE cells can be delivered in the subretinal space in an ex vivo model

In order to assess the feasibility of delivering cells to the subretinal space with a microneedle, we utilized a closed vitrectomy model to assess whether RPE cells can be delivered in the same manner as previously described. Twenty-five freshly harvested 3-6 month old pig eyes were used. We found in all cases, it was possible to inject iPS RPE cells into the subretinal space without any surgical impairment. A retinal bleb was formed consistently and there were no needle blockages in any of the procedures. No reflux of the cell solution was noted following needle removal from the retinotomy site when using the 49G microneedle but occasionally reflux was evident with the 41G and rarely the 43G microneedles (Supplementary information – S4, S5, and S6). Spectral domain optical coherence tomography imaging (SD-OCT) performed immediately after the procedure, confirmed the presence of a subretinal bleb (Fig. 7 F-H), however, it was not possible to determine cellular reflux, which was examined with a high concentration of cells injected purposely at the pre-retinal space. In some eyes, it was possible to detect fluorescent microbeads, however this was inconsistent and a positive control with a high concentration of fluorescent microbeads injected into the pre-retina space was not visible. This was validated by also attempting the same imaging with swept source OCT, which could not resolve the injected cells. In order to further assess the feasibility of minimally invasive surgery, we injected the iPSC RPE cell suspension in eyes without performing vitrectomy, finding that this was feasible and free from any blockage or intraoperative complication (Fig. I & J). Histological examination demonstrated persistent retinal detachment, expected in a cadaveric eye without functional or damaged RPE, and the presence of iPSC RPE cells attached to the outer neurosensory retina confirmed with CD147 antibody (Fig. 8).

**Figure 7:**
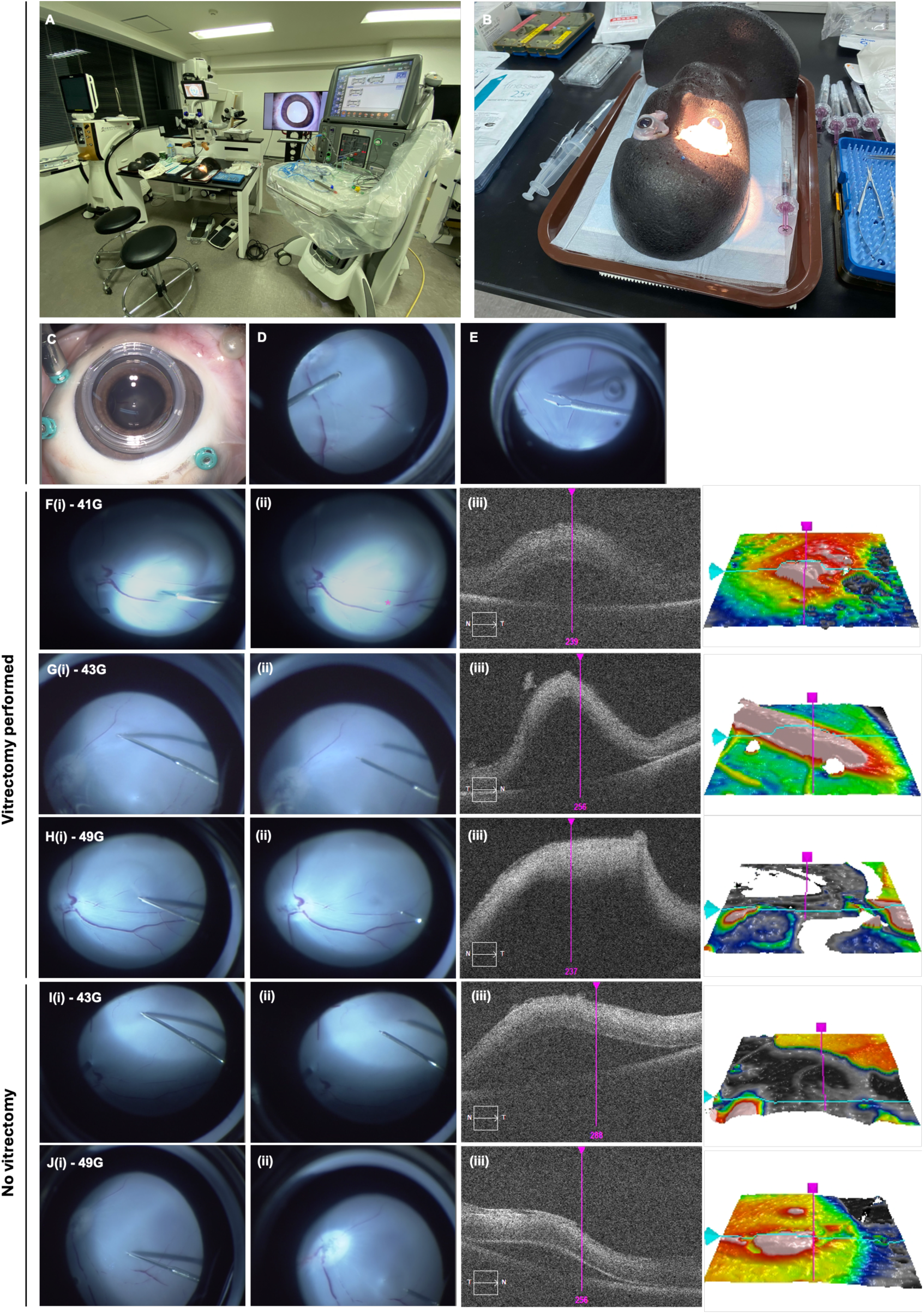
Ex vivo cadaveric porcine subretinal surgical delivery of iPSC RPE cells. (A) Ex vivo porcine eye vitrectomy set up with (B) eyes secured to a phantom head for surgery. Intraoperative images of (C) 3 port vitrectomy, (D) end-gripping forceps used to peel the posterior hyaloid/internal limiting membrane when required, and (E) subretinal injection of iPSC RPE cells and 1:100 fluorescent microbeads solution. Ex vivo spectral domain OCT images of subretinal blebs created by injecting (F) 41G microneedle, (G) 43G microneedle, and (H) 49G microneedle following vitrectomy and without…., respectively, following vitrectomy. Eyes where no vitrectomy was performed and only the subretinal injection of iPSC RPE cells, it was possible to form a subretinal bleb using the (I) 43G needle and (J) 49G microneedle. Due to the inability of the soft malleable tip of the 41G needle to penetrate vitreous, a vitrectomy was necessary to create a subretinal bleb.

**Figure 8:**
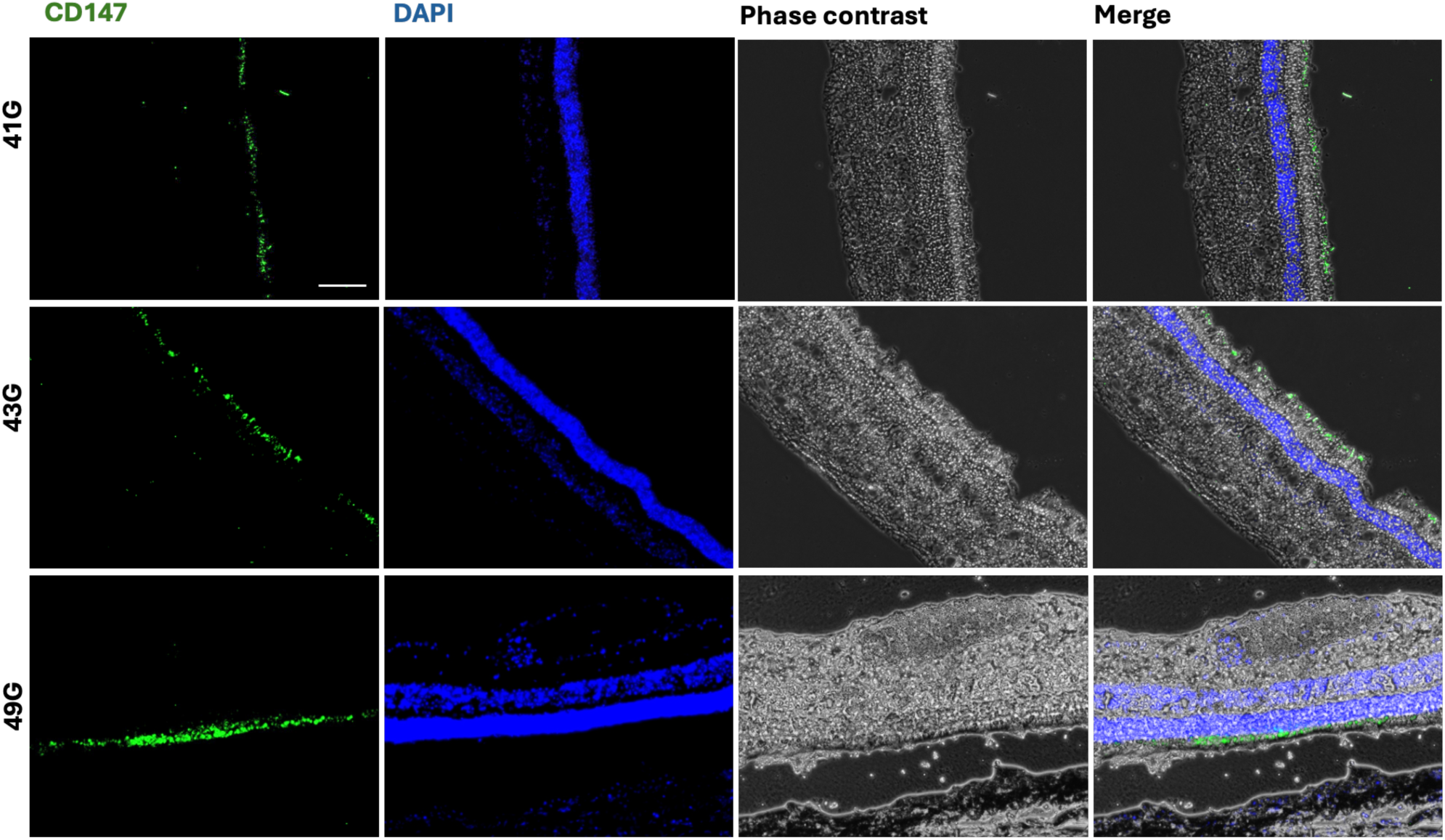
Ex vivo porcine eye iPSC RPE cells labelled with CD147 (green) implanted in the subretinal space. Imagees for 41G and 43G are of fully detached retina, while 49G images show shallow retinal detachment. In all images iPRSC RPE cells have adhered or engrafted into the cadaveric porcine outer retina, confirming feasibility of microneedle cell delivery. Scale bar in top left image represents 100µm and applies to all images in the panel.

## Discussion

Minimally invasive surgery has progressed over the decades along with manufacturing technologies that permit less traumatic and safer ways of delivering therapeutic agents. In dermatology, microneedles hold great promise in delivering a range of treatments including regenerative therapies, vaccinations, hormones and immunotherapies. Here, we demonstrate for the first time, that a microneedle with an estimated bore of 49G can be used to surgically implant iPSC RPE derived cells by using in vitro and ex vivo models. Notably, we utilized surgical instrumentation, including the microneedles, that are currently used in clinical practice internationally. Further, the iPSC RPE cells have been validated as safe for human use, making the results of this work highly translatable in the field of retinal surgery and potentially in any clinical field delivering cell therapy at the microscale. To our knowledge, this is the smallest scale from which cells can potentially be surgically delivered for human therapy.

Microfabrication technologies have led to the ability to manufacture solid microneedles with tips as small as 1µm, which have been shown to be painless when applied to human skin.^20^ Hollow silicone-based microneedles with bores as small as 75 µm have been fabricated and shown to effectively deliver melanocytes, keratinocytes and epidermal cells to skin.^21^ Solid microneedles designed for delivery of cell therapies are fabricated with a range of materials including polymethyl methacrylate (PMMA)^22^, hydrogels^23,24^, frozen phosphate buffer solution and dimethylesulfoxide^25^, and poly(lactic co-glycolic) acid^26^ and can be designed to be porous or coated to harbor cells for example. These approaches are promising in a multitude of fields where minimally invasive cell transplantation can be beneficial and longer lasting. In ophthalmic surgery, there are numerous hollow microneedles that have been employed in established and emerging therapies. Recent clinical applications of a range of short 30G needles designed specifically for suprachoroidal steroid therapy have been developed successfully and marketed as Xipere®.^27^ Glass hollow microneedles have been explored for intrascleral and transscleral therapeutic delivery in various experimental models.^28,29^Essentially, these microneedles are short in length with a diameter in the region of 30G and less than 1000 µm in length, for example 7—to 750 µm 30G needles have been used in hydrogel delivery to the suprachoroidal space and collegenase.^30,31^ We were therefore interested to investigate the effect of bore size in the smallest microneedles currently available for human surgery, to determine the feasibility of minimally invasive cell delivery.

A previous study by Wilson et al. investigated the optimal needle size by assessing immediate cell viability after repeated handheld pipetting of 10µL of suspensions of iPSC RPE cells using trypan blue staining.^32^ They found a positive correlation between a smaller needle gauge and percentage cell viability, notably with the 38G and 41G canulae a viability of 15-20% was found, while their control and 30G needle elicited a viability of around 90%. In comparison we found an average of 92%, 96%, 79% and 83% cell viability with control, 41G, 43G and 49G needles, suggesting a number of reasons for such a difference in findings, particularly with the 41G cannula used in both studies. In the present study, a Constellation VFC system was used to inject the iPSC RPE cells through a MicroDose 1mL syringe using consistent injection pressures throughout all repeat experiments. Wilson et al. utilized a 250µm borosilicate glass syringe (Hamilton), commonly used in animal experimental studies, to manually inject cells with inevitable variability in the applied pressure across the experiment. Such variation in injection pressure results in inconsistent fluidic currents within the needle, which may result in variable degrees of cell stress and cell shear. This may represent some of the reasons for the difference in cell viability analysis between these experiments, however other factors such as the cell wall elasticity between RPE cell lines, as shown between different stem cell types, may play a role in this.^33^ Further, our experiments have been carried out three times at separate time points using two different cell lines, a primary and stem cell derived cell lines, increasing the robustness of the investigation. The concerns Wilson et al. raise regarding cellular debris and a high percentage of dead cells being delivered using microneedles in the subretinal space appear unlikely given the results of our study and prior studies using 41G subretinal needles to successfully deliver stem cell derived RPE cells to both animals and humans.^6,7,14^ Altogether, the need to use a 30G needle to deliver RPE cells into the subretinal space as described by Wilson appears unnecessary, where we should that it is feasible to use a minimally invasive microneedle instead.

Previous studies examining the effect of needle bore size on cell survival, function and phenotype have not found any deleterious effects. For example, Mamidi et al. compared injecting mesenchymal stem cells (MSCs) through 24G, 25G and 26G needles into culture dishes, finding that there was no effect on cell morphology, viability, differentiation potential and in vivo migration.^34^ Agashi et al. utilized a 22G, 25G and 26G needle to inject murine MSCs at different flow rates, finding that smaller bore size as well as longer lengths of time the cells are kept inside the syringe prior to injection, negatively affected cell viability.^35^ Similarly Lang et al. examined the effect on equine MSC viability when injecting with 20G, 22G, 23G, and 25G needles, finding that larger bore needles improve cell viability.^36^ The smallest bore microneedles investigated for use in cell therapy was 34G. Amer et al. compared 20mm long 30G and 34G needles when injecting human MSCs, finding that the smaller gauge needle and slower rates of infusion had a negative impact on cell viability.^37^ They found that at the lowest flow rate of 10μL/min, significant apoptic cell death occurred in both 30G and 34G needles, with 20, 50 and 150μL/min flow rates causing non-significant changes in apoptic cell death. However, at the highest flow rate of 300μL/min, the 34G but not the 30G needle was associated with significantly higher apoptic cell death. Our study purposely did not employ an infusion pump because the goal was to investigate minimally invasive cell delivery using surgical equipment in current clinical practice. While we did not examine specific injection pressures, we utilized varying cell densities to clarify whether small gauge needles cause significant cellular shear and needle occlusion. We demonstrate that RPE concentrations of 3.0 x 10^4^, 6.0 x 10^4^, 2.5 x 10^5^ cells can be injected without any significant loss in viability, morphology or function. Further, we estimate the maximum shear stress of each needle, finding variability in average predicted shear stress and hypothesis that central flow with the least shear may account for the lack of difference in viability between the microneedles and control. Others have reported that low shears of 15Dyn/cm^3^ can affect stem cell differentiation^38^ and therefore shear forces may influence different cell types in different manners. Herein, we report shear as high as 2.36 x 10^3^ Dyn/cm^3^ with no significant loss of viability compared to control, indicating that fluid dynamics through microneedles may not behave as expected when injecting cell suspensions. Cell aggregations through small bore needles is a concern when delivering cell therapies^39^, however, we found that microneedles did occlude at any stage throughout these experiments.

As previously described, cell viability assays are important assessments of the effect of the needle characteristics on cell therapy.^37^ However, shear forces beyond physiological levels may activate signaling cascades or apoptic events that can lead to delayed cell death not detectable by trypan blue exclusion assay for example. We therefore monitored cell viability with SYTOX green over 2 weeks of cell culture to mitigate these effects and went on to examine morphology and function. Altogether, we demonstrate that RPE cells can robustly attach, proliferate and function after passage through microneedles as small as 49G. As the manufacture of 43G and 49G microneedles is a multi-step process requiring multiple specialist manufacturers to fabricate and then assemble these products, there is a limited supply at any given time. This also results in a relatively high cost, making it challenging to design an experiment that can employ an ideal range of experimental techniques commonly used in the field with widely available small gauge needles such as the aforementioned 25G, 26G, 27G, 30G, 34G needles. We therefore were unable to undertake fluorescent cell sorting to measure cell size but appreciate this would be useful to carry out in future work. A strength of our study is that we used both primary and stem cell derived RPE cells, showing similar results and increasing the robustness of our results.

In order to gain insight into shear stress that may be elicited from passage through a small needle bore, we examined the expression of F-actin to understand any cytoskeletal abnormalities that may occur. Cucina et al. previously showed that vascular endothelial cells undergo cytoskeletal re-modelling when undergoing laminar flow where F-actin loses its array of microfilament bundles of stress fibres and takes on a diffuse pattern.^40^ RPE cells are known to express F-actin as lateral circumferential pattern in physiological conditions, however, it has been shown this becomes disorganized in stem cell derived RPE cells that have low phagocytic activity.^41^ In our experiments, we found that there were no adverse changes to F-actin distribution in both primary and stem cell derived RPE cells after ejection from any of the microneedles tested. Despite this, due to experimental constraints (i.e. cell number), we did not carry out a time course on F-actin expression, which could mean that any cells with significant shear stress may have detached and washed away after the required media changes. This is important to appreciate as cellular debris is thought to act as a detrimental sources of cytokines and inflammatory mediators that may compromise any engrafted cells.

The ex vivo porcine data of subretinal RPE transplantation provided herein, provides useful data that can be used to determine the feasibility of minimally invasive transplantation of iPSC RPE cell transplantation. A disadvantage of this is that cellular reflux following cell suspension injection into the subretinal space cannot be assessed accurately, nor can the formation of epiretinal membrane. We found that SD-OCT was unable to resolve the microneedles retinal wound tract easily and this may be due to resolution limitations met with the possibility of self-sealing retinal wound formation with the 43g and 49G microneedles. Further, when testing whether human iPSC RPE cells can be detected by OCT when implanted in a pig eye, both SD-OCT and swept source OCT was unable to identify the cells. Histological analysis did not permit the identification of the retinal wounds and therefore it would be beneficial to examine this in a live animal model. An important advantage of the current study is that the use of live animals was avoided to obtain feasibility data, in keeping with the widely accepted 3R principles.^42^ Finally, we were able to demonstrate that subretinal transplantation may be feasible without a vitrectomy, akin to modern choroidal biopsy techniques.^43^ This may therefore be developed in the future as a minimally invasive surgical technique for retinal cell therapy.

## Conclusions

Herein, we explore the feasibility of minimally invasive cell transplantation surgery, finding that microneedles can deliver both primary and stem cell derived cells without causing significant loss in cell viability, cell morphology or cell function. The use of microneedles can potentially abrogate the need for a vitrectomy to deliver cells to the subretinal space, indicating a less invasive and safer procedure may be possible in the future. Further, it may be possible to deliver current and emerging therapies such as anti-VEGF antibodies and gene therapy using microneedles. As such, these findings may represent a future paradigm shift in minimally invasive retinal therapeutics.

## Acknowledgements

We are most grateful to Mr Hashimoto Koichi and all his colleagues at the Alcon Surgical Training Center, for all their expertise and allowing us to use their facilities for vitrectomy surgery. We are very grateful to Naoki Yoshihara for his support and expertise in taking all electron microscopy images at the Instrumental Analysis Center, Research Initiatives and Promotions Organization, Yokohama National University. We thank all members at the Department of Ophthalmology and Micro-technology, Yokohama City University.

## Funding

J.C. acknowledges funding from the Keeler Scholarship and T.F.C. Frost Charitable Trust.

## Author Contributions

This project was conceived and initially developed by J.C., S.K. and K.K. Benchtop experiments and ex vivo porcine experiments designed by J.C., S.K., I.H. and performed by J.C and I.H. Data collection and analysis were carried out by J.C. and I.H. Manuscript draft preparation and finalization undertaken by all authors.

## Competing interests

K.K. holds a patent for the 45G/49G needle described in this work (Patent No. 5777074).

## Supplementary Information

**S1:**
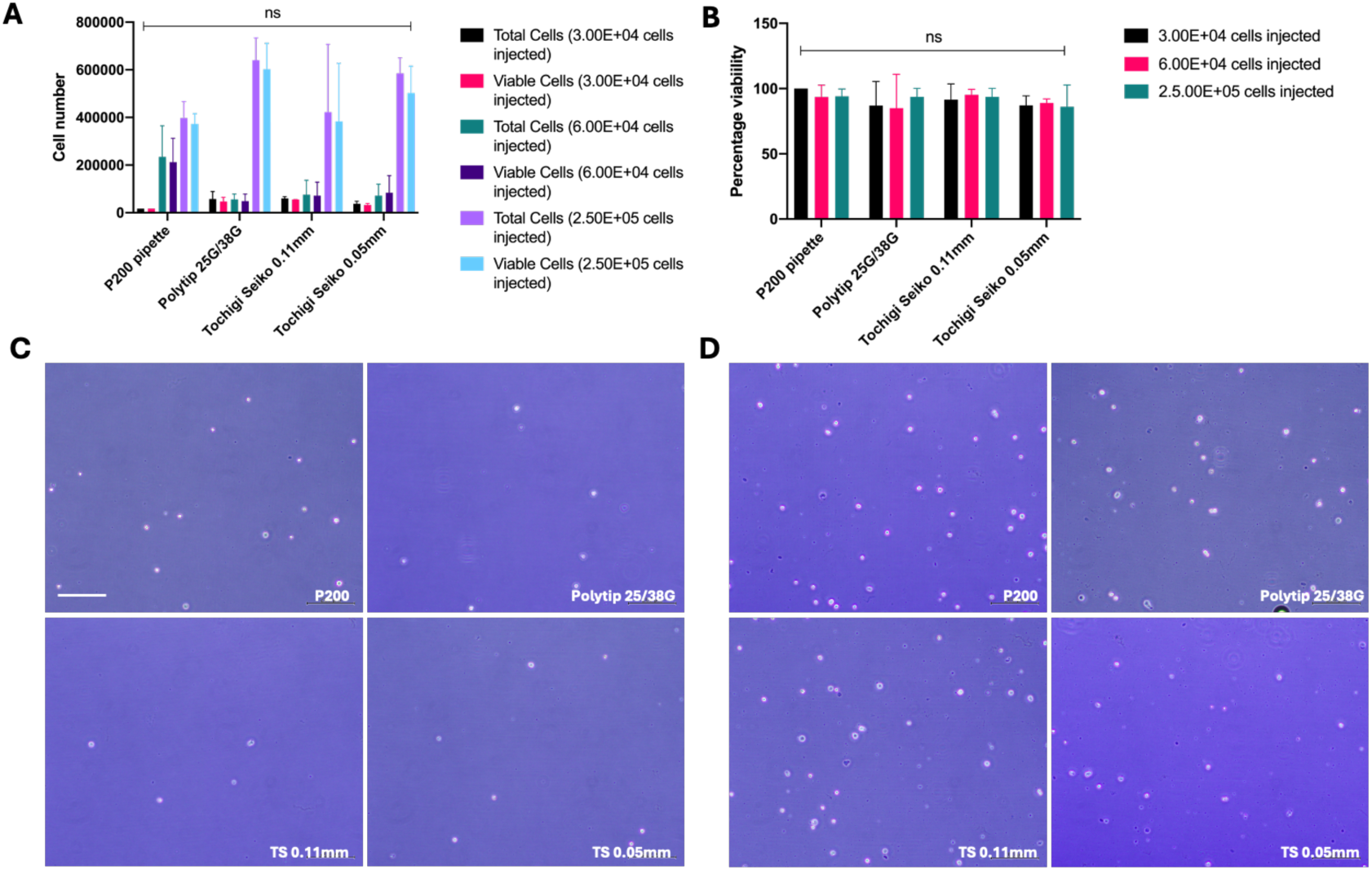
Cellularity and viability of primary RPE cells injected from microneedles. (A) Total and viable cells injected from control (P200) and different microneedles as calculated from trypan blue exclusion assays using different concentrations of cells. (B) Percentage cell viability for control and microneedles with different cell concentrations. (C) Trypan blue exclusion assay image under bright field microscopy for primary RPE cells injected at a density of 6 x 10^4^ in 150µL, scale bar represents 500µm and applies to all images in (C) and (D). (D) Trypan blue exclusion assay image under bright field microscopy for primary RPE cells injected at a density of 2.5 x 10^5^ in 150µL.

**S2:**
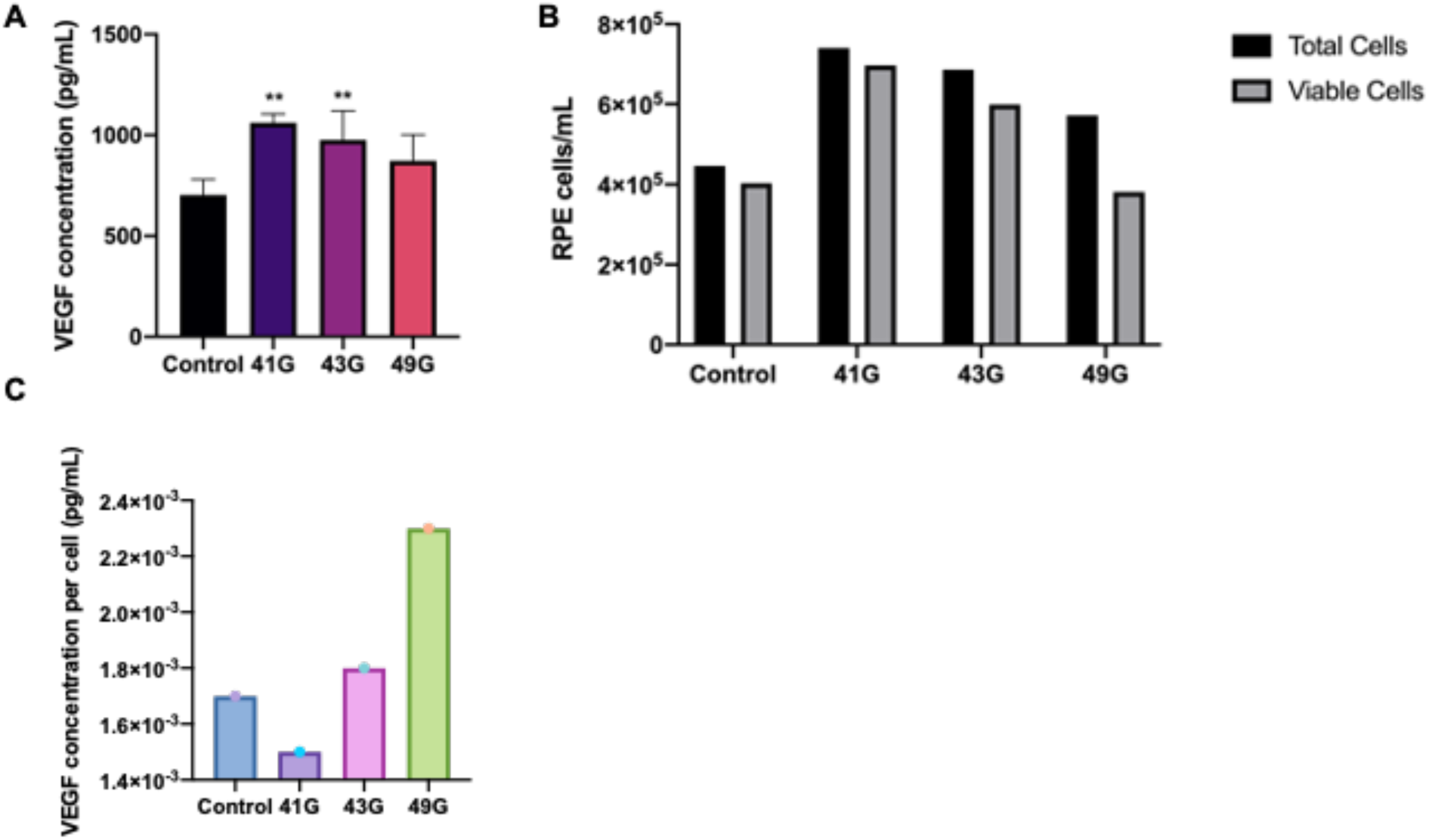
(A) Primary RPE VEGF secretion measured by ELISA. ** - p < 0.001. (B) Total and viable RPE cells measured for experiment media for VEGF analysis derived from. (C) Controlling for cell number, the amount of VEGF secreted per cell was within a range of 1.0 x 10^-3^pg/mL between all conditions.

**S3:**
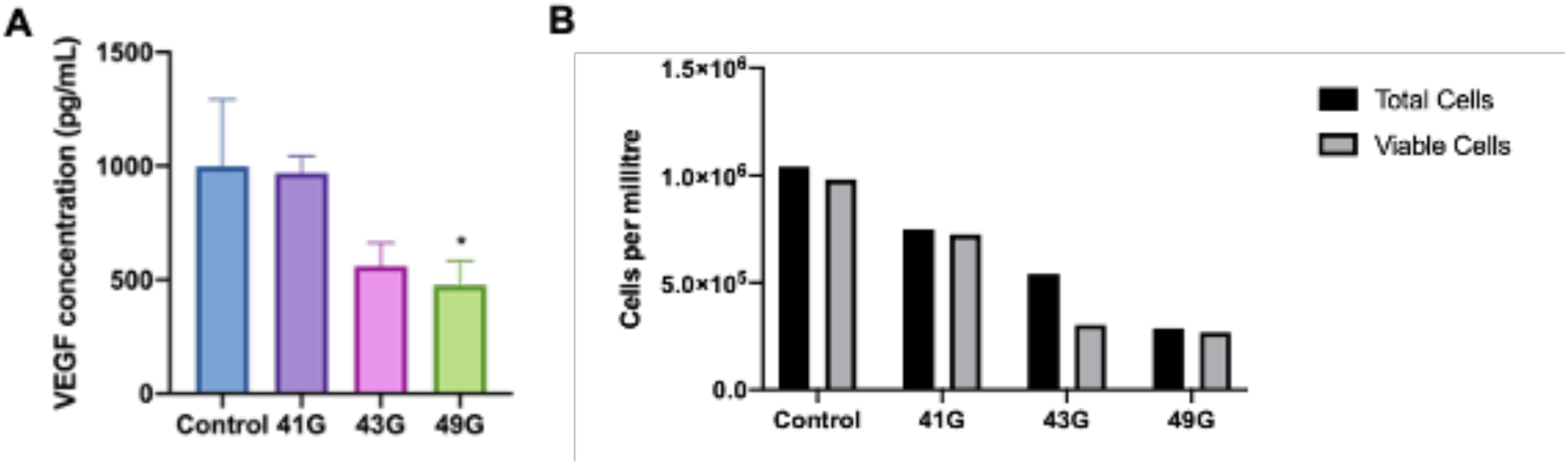
(A) iPSC RPE VEGF secretion measured by ELISA. * - p < 0.05. (B) Cell counts of iPSC RPE cells of the corresponding experiment from which cell culture media was taken and analyzed for VEGF by ELISA.

**S4:** Intra-operative video of subretinal implantation of iPSC RPE and fluorescent microbeads (1:100) suspension using a 41G needle with a 25G three port vitrectomy approach with a core and peripheral vitrectomy. A plume of suspension can be seen when the needle is withdrawn from the retina.

**S5:** Intra-operative video of subretinal implantation of iPSC RPE and fluorescent microbeads (1:100) suspension using a 43G needle with a 25G three port vitrectomy approach without vitrectomy. A plume of suspension can be seen as the needle is withdrawn.

**S6:** Intra-operative video of subretinal implantation of iPSC RPE and fluorescent microbeads (1:100) suspension using a 49G needle with a 25G three port vitrectomy approach without vitrectomy. No suspension can be seen when the needle is withdrawn.

